# Divergent impact of grazing on plant communities of created wetlands with varying hydrology and antecedent land use

**DOI:** 10.1101/2020.06.15.153155

**Authors:** Kimberly A. Lodge, Anna Christina Tyler

## Abstract

Destruction of natural wetlands has warranted the creation of wetlands to mitigate the reduction of valuable ecosystem functions and services. However, the complex interactions between key drivers of wetland community structure – hydrology, nutrient availability and herbivory – makes creation of functional wetland replacements challenging. We examined interactions among these drivers, and their impacts on plant communities and soil characteristics in two created wetlands with different hydrology and land use histories: a shallow emergent marsh on a previous gravel depository and a seasonally flooded marsh on a former cattle pasture. In paired plots open to or protected from large wetland grazers we observed seasonal variation in grazing impacts on plant communities and an increase in effect size over time. At the permanently flooded marsh with high grazing waterfowl densities and low nutrients, open plots had significantly reduced plant growth and diversity, and an increase in submerged vegetation. In contrast, grazer density was lower and nutrients were higher in the seasonally flooded marsh, where grazer access enhanced plant diversity and reduced invasive plant cover. These results suggest the possibility of long-term grazer-induced shifts in community composition and delivery of key ecosystem services in young, vulnerable created wetlands. To improve created wetland design and function, we suggest that in addition to hydrologic conditions, the impact of prior land use on present nutrient availability be considered. Further, enhanced heterogeneity of spatial and bathymetric structure can provide conditions for diverse plant communities and balanced habitat use by wetland grazers.

## Introduction

The abiotic and biotic structures and processes – transformations of matter and energy – that take place within wetland ecosystems result in the production of ecologically, economically, and culturally valuable natural resources (Costanza et al. 1997; Costanza et al. 2014; Georgiou and Turner 2012). A key aspect of wetland function and habitat provision is the presence of emergent macrophyte communities which influence biogeochemical processes and, in turn, are shaped by both biotic and abiotic factors (Rejmankova 2011; Tanner and Headley 2011; Thomaz and Cunha 2010). The primary factors that drive plant diversity and ecosystem function in natural wetlands – hydrology, nutrient availability, and herbivory – interact in complex and dynamic ways that are not fully understood, complicating the planning and design of projects related to wetland creation and restoration (Bakker et al. 2013; Bakker et al. 2016; Engels and Jensen 2010). Created wetlands are prone to colonization by invasive plants and may have lower species richness without active management and planting native species; this leads to the development of alternative community structure and successional trajectories (LePage 2011; Matthews and Spyreas 2010; Stefanik and Mitsch 2012). In order to enhance the success of wetland creation and restoration attempts, a better understanding of abiotic and biotic controls on wetland structure and function is required.

Hydrology is the ultimate driver of wetland structure and function, and variations in surface water depth and duration of saturation regulate soil and water chemistry by determining oxygen availability and soil redox status; this indirectly controls nutrient availability for competing plants and microorganisms (Jahangir et al. 2016; Song et al. 2012), and directly controls the number, type, and distribution of individual plant species (Gurnell et al. 2012; Zhang et al. 2012). The resulting plant communities provide food and habitat for wetland herbivores, such as waterfowl, which choose their habitats based on water depth and resource availability (Bino et al. 2014; Pickens and King 2014). When wetlands are created by altering hydrology of land previously used for other purposes, decisions made during the planning and construction can alter habitat suitability for waterfowl and other grazers, changing potential grazer influence on wetland function. The consequences of these alterations in created wetlands is important to more fully understand.

Aquatic herbivores carry out key top-down controls on community dynamics through the selection of plant species based on the nutrient content, palatability, or physical stem structure (Duarte et al. 2014; Gutbrodt et al. 2012; Morrison and Haye 2012). At high densities and intense grazing, herbivores can significantly reduce above- and below-ground biomass of preferred plants; this drives plant competition as unpalatable species gain a competitive advantage and ultimately lead to long-term reductions in plant community diversity (Baldassarre et al. 2006; Bertness et al. 2014; He and Silliman 2016; Wood et al. 2012). Foraging behaviors also have strong implications for invasive species success in created wetlands; native plant species are often preferred by native grazers, and rapid clonal growth of invasive species prevents significant control by herbivores (Grosholz et al. 2009; Li et al. 2014). Created wetlands may be especially vulnerable to grazer-induced shifts in community composition, because their plant communities are young and less resilient than mature, natural wetlands. This vulnerability may be exacerbated by the massive increases in grazing waterfowl populations observed in recent decades (Ankney 1996).

Herbivores may also alter soil dynamics through direct disruptions to the soil layers during root or rhizome removal, and through indirect alterations to nutrient pools in the soil (Kotanen and Abraham 2013; Pulido et al. 2016). The deposition of nutrient-rich feces while foraging can increase both nitrogen and phosphorus levels (Mallin et al. 2016; Telesford-Checkley et al. 2017); however, removal of nutrient-rich plant material before moving on to another area can decrease local nutrient availability (Kitchell et al. 1999; Metcalfe et al. 2014). Also, reallocation of resources by damaged plants, and increased nutrient uptake for recovery growth may lead to decreased nutrient pools and root exudation of labile carbon (Gao et al. 2008; Piñeiro et al. 2010). Continued reductions in plant biomass may further decrease carbon availability by lowering litter input during senescence (Bai et al. 2012; Medina-Roldán 2012; Yan et al. 2013). This could pose further problems in the development of communities in created wetlands, which often have significantly lower soil organic matter than comparable natural wetlands (e.g., Campbell et al. 2002; Fennessy et al. 2008).

The impact of wetland herbivores on plant communities can be significant, but interactions with nutrient availability and hydrology likely contribute to the community-level response and resilience to disturbance. Resource limitation in nutrient-poor ecosystems may prevent plant regrowth after grazing; however, grazing in nutrient-rich ecosystems may actually facilitate competition and diversity (Fornoni 2011; Proulx & Muzumder 1998). Considering the constant competition for resources between plants and microorganisms, the level of nutrients within the system becomes important to combat detrimental impacts of grazers. Unique to created wetlands is the legacy of prior land use on present nutrient availability; this link is not clearly defined, but may have cascading impacts on the ability of plant communities to recover from grazing events (Foster et al. 2003). These impacts can be especially pronounced during migration periods or at over-wintering grounds when grazer populations are at their highest and may result in limiting the re-establishment of plants in subsequent growing seasons (Chaichana et al. 2011; Guillaume et al. 2014; Bakker et al. 2018).

The overarching goal of our study was to evaluate the interactions between hydrology, nutrient availability, and herbivory in created wetlands in order to inform the design and management of similar systems before and after construction. These interactions were evaluated in two created, emergent freshwater wetlands with different prior land use histories: wet and low nutrients, dry and high nutrients. While these systems don’t allow a full factorial analysis of driving forces, the contrasting nature of these systems allows a unique opportunity for comparison of the controls on wetland structure-function relationships in created wetlands. We hypothesized that: 1) grazing pressure will be higher in created wetlands that are permanently flooded, and will shift seasonally in time with migration cycles, 2) the presence of grazers will decrease both plant growth and diversity, with a greater effect where grazing pressure is high, and 3) the removal of plant matter by grazing will decrease soil nutrient pools and organic matter.

## Methods

### Study sites

This study was conducted from 2014 to 2016 in two created wetlands at High Acres Nature Area (HANA) in Perinton, New York, USA, both of which are owned and managed by Waste Management of New York and New England, LLC. Both wetlands were created as mitigation projects associated with a landfill expansion. Area 1 North (A1N; 43°05’33.37”N, 77°23’10.34”W) was created in 2009 as a 1.87 ha shallow marsh. The site served as a gravel-mining depository until approximately the mid-1960s, before being abandoned and left fallow. Prior to mining, the area was used for agricultural purposes. At the time of construction and in subsequent years, native emergent species – *Sagittaria latifolia, Alisma plantago-aquatica,* and *Polygonum* spp. – were planted and were the dominant species at the start of this study. However, immediately following construction, invasive *Typha latifolia* and *Typha angustifolia* colonized the majority of the site, leading to extensive control executed via manual cutting, pulling, and herbicide applications (glyphosate) between 2010 and 2016 that significantly reduced the cover of *Typha* spp. Following the initiation of this study, no intentional invasive plant control was conducted in the vicinity of the treatment plots. The hydrology of A1N is driven by subsurface flow from an adjacent abandoned quarry pond and precipitation; a culvert at the southeast corner allows water control and maintenance of standing water year-round.

Area 3 (A3; 43°05’16.14”N, 77°22’15.06”W) was a cattle pasture prior to construction of approximately 1.63 ha of shrub/scrub and shallow marsh area in 2012. Groundwater flow from an adjacent hillside and precipitation are the dominant hydrologic drivers at this site; there was no water control capacity during the study period. At the time of construction and in subsequent years, native shrub and emergent wetland species were planted throughout and *Typha* spp., *A. plantago-aquatica,* and *Polygonum* spp. were dominant at the initiation of this study. *Typha* spp. seed heads were cut each summer and plants were sprayed with herbicide (glyphosate) each fall since 2013, avoiding experimental plots.

### Characterization of grazing pressure

Abundance of large wetland herbivores was quantified from September 2015 to September 2016 by the authors and trained volunteers. We recorded the number and species of grazers present (including tracks and houses/nests), their behavior (foraging, swimming, nesting, etc.), date, and time of day, during all visits to the two wetlands. The frequency of observation varied between the two wetlands, but was sufficient to demonstrate differences in grazer identity and density between sites. Results were converted to grazer density (area calculated using ArcGIS mapping software) across seasons (spring, summer, fall, and winter; Marklund et al. 1992).

### Experimental design: herbivore exclusion

In June 2014, 16 pairs of 1 m^2^ caged (herbivore exclusion) and uncaged control (open to herbivores) plots arranged in four blocks of four pairs, were established randomly at each site (64 total plots). Paired caged and uncaged plots were 1 m apart and at least 3 m from another pair. Caged plots were set up by wrapping galvanized hardware cloth (1.27 cm mesh, 1.22 m tall) around four polyvinyl chloride pipes (PVC); uncaged plots were marked with PVC pipes only. In May 2015, one additional three-sided caged plot was established at each block to evaluate potential cage effects (8 total cage-control plots).

### Hydrologic conditions and soil nutrient availability

Hydrologic conditions were evaluated by averaging surface water depths from 3 points in every plot in spring (May), early-summer (June), mid-summer (July), and fall (September) of each year. Three soil cores (2.5 cm diameter x 10 cm deep) were extracted from each plot with an auger in September 2014, 2015, and 2016 and in May 2016, and subdivided for organic matter and nutrient analysis. Soil organic matter content was determined using the loss on combustion method (Heiri et al. 2001). Inorganic nitrogen was extracted by shaking wet soil with 2M potassium chloride (Keeney and Nelson 1982). Ammonium was analyzed using the phenol-hypochlorite method and a Shimadzu 1800 spectrophotometer (Solorzano 1969). Nitrate+nitrite was measured with the cadmium reduction method and a Lachat Quikchem 8500 autoanalyzer (Lachat 2003). Total inorganic nitrogen (TIN) was calculated by summing extractable ammonium and nitrate+nitrite. Total phosphorus (TP; May and September 2016 only) was extracted from soil samples by adding magnesium nitrate to soil dried at 60°C, ashing in a 550°C oven for two hours, and dissolving in sulfuric acid before analysis using the ammonium molybdate method (Murphy & Riley 1962).

### Plant cover, diversity and belowground biomass

Vegetation measurements were collected at three to four time points during the growing season in all plots starting in June 2014 and ending in September 2016. Percent cover of each species was estimated by at least 2 observers per plot and averaged to eliminate observer bias. Plant diversity was evaluated using species richness (*S*) and the Shannon-Weiner Diversity Index (*H*’). Plant stem density and height (three tallest stems of each species) were measured in May, June, July and September of 2016 for all species. Belowground biomass was measured in September 2016. One soil core (10 cm diameter x 25 cm depth) was collected from each plot using an auger, washed through a 1 mm mesh sieve to remove soil particles, dried (60°C) and weighed (Evers et al. 1998).

### Statistical analyses

All statistical analyses were completed using JMP 13 Pro statistical software. Prior to selection of statistical analysis method, each dataset was checked for normality and homogeneity of variance. Grazer density was analyzed using a one-way Kruskal-Wallis test to compare total average individuals per hectare between sites within seasons (i.e. spring A1N vs spring A3, etc.).

To characterize between site differences, we made comparisons of control plots only for organic matter, total inorganic nitrogen, and total phosphorus using a one-way analysis of variance (ANOVA). We also used a full-factorial three-way ANOVA to compare intra-site differences between these variables with treatment (caged/uncaged), season (spring, fall), and year (2014-2016), when applicable, as fixed factors.

Statistical analyses for effects of grazing on stem height, stem density, and percent cover for individual species were conducted only for species with a percent cover ≥5% in at least five plots across the growing season. Calculation and comparison of total plant cover and diversity indices (*S* and *H*’) included minor species. All *Polygonum* species and *Typha* species were grouped together for plant height analyses. We made between site comparisons of total plant cover, *S*, and *H*’ using a one-way ANOVA, only including uncaged control plots in this analysis. Using a full factorial three-way ANOVA we compared differences within sites for all other plant variables with treatment (caged/uncaged), season (spring, early summer, mid-summer, fall), and year (2014-2016), when applicable, as fixed factors. Invasive species cover was evaluated for A3 only because of the low cover (typically <1 %) in A1N in any one season. For all variables, when significant differences were found, a Tukey’s HSD post hoc analysis was used to elucidate differences among treatments.

## Results

### Hydrology

A1 N was flooded throughout the year, whereas A3 was seasonally flooded and was typically dry by early July. The average water depth between May and September was consistently two or more times deeper in A1 N than A3 (Fig. 1). A drought in 2016 decreased average water depths in both wetlands, and resulted in A3 completely drying by mid-June.

**Fig. 1.**
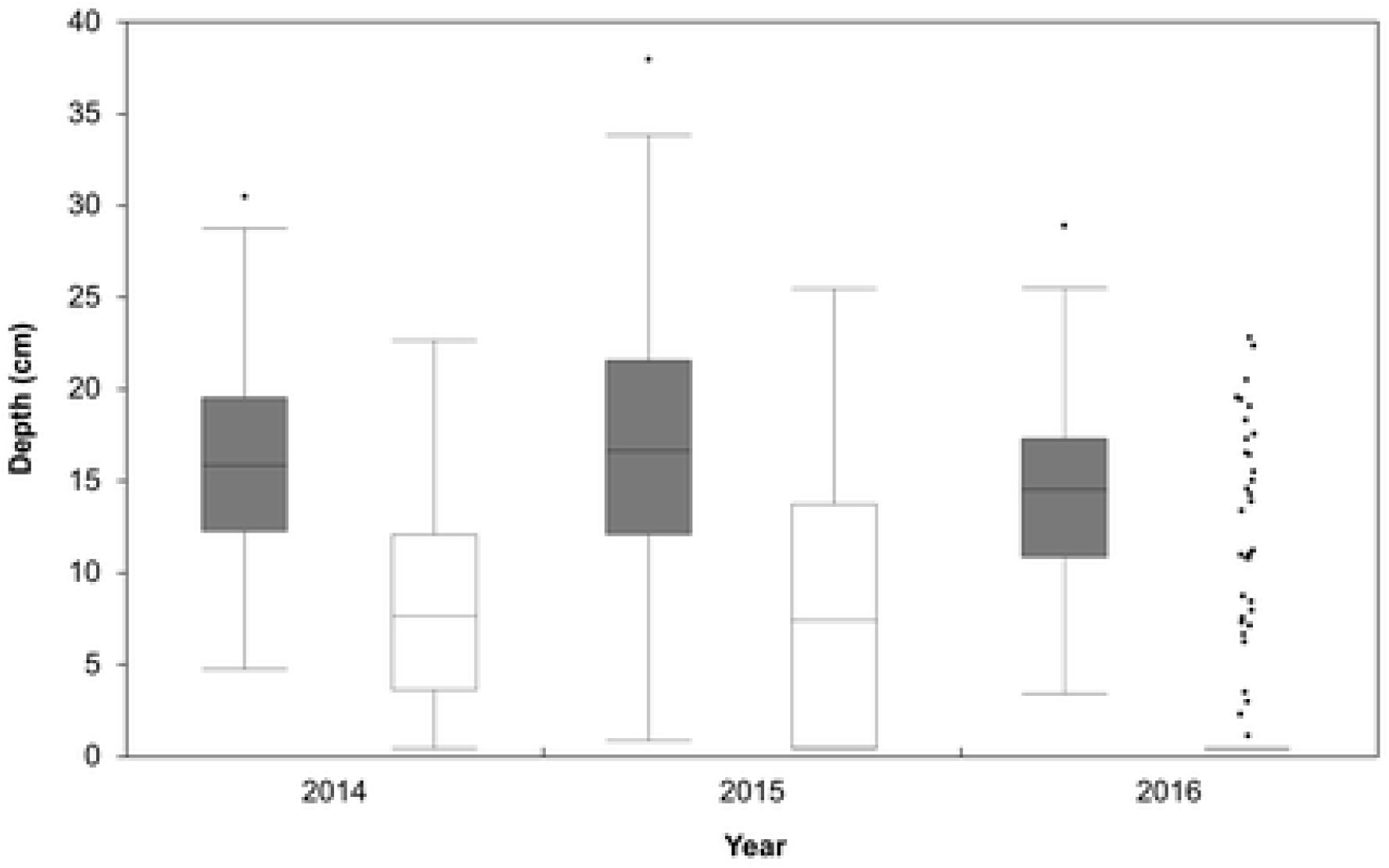
Boxplots of water depth across the growing season (May-September) in Area 1 North (gray) and Area 3 (white) from 2014-2016. New York State experienced a drought beginning in June 2016.

### Grazing Pressure

At both sites we observed Canada goose (*Branta canadensis*), ducks (*Anas* spp.), whitetail deer (*Odocoileus virginianus*), North American beaver (*Castor canadensis*), and the common muskrat (*Ondatra zibethicus*). Waterfowl comprised the majority of grazer abundance at both sites: 99-100% and 66-100% of grazers were waterfowl at A1N and A3, respectively. Overall grazer density was significantly greater in A1 N than A3 (41.2 ± 7.4 and 4.9 ± 1.5 individuals ha^-1^, respectively; χ^2^ =41.9, p <0.0001; Fig. 2), but the relative difference varied across seasons. Grazer density was roughly 90 (summer) and 8 (fall, peak) times higher in A1 N than A3 (χ^2^ =18.2 and 26.4, respectively, p <0.0001), but were similar in spring and winter (χ^2^ <1, p =0.95 and χ^2^ =1.3, p =0.25, respectively).

**Fig. 2.**
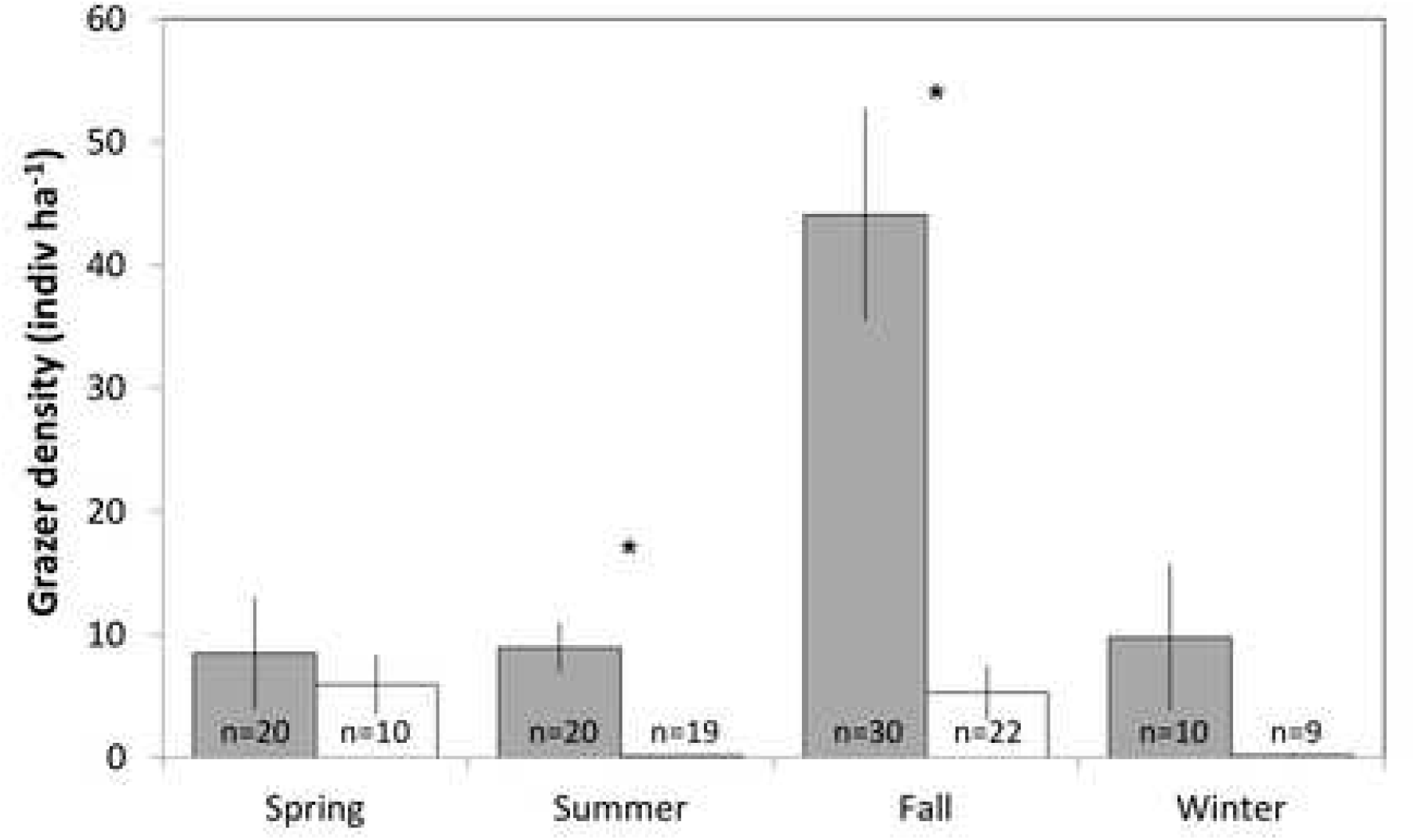
Mean ± SE large grazer density observed at Area 1 North (gray) and Area 3 (white) wetlands between September 2015 and September 2016 (spring = March-May, summer = June-Aug, fall = Sept-Nov, winter = Dec-Feb). Text values on bars are the number of individual observations per season. * indicates significant difference between sites within a season (p <0.0001).

### Nutrient availability

Soil nutrients and organic matter in control plots were consistently higher in A3 relative to A1 N (Table S1): organic matter content (OM) was 1.5 times greater (13.4 ± 0.5 versus 7.5 ± 0.4 %; p <0.0001); total inorganic nitrogen (TIN) was 3 times greater (17.1 ± 3.4 versus 6.2 ± 1.8 mg/kg; p <0.0001); total phosphorus (TP) was 1.5 times greater (1002.7 ± 53.2 versus 704.3 ± 28.0 mg/kg; p <0.0001), respectively. In A1N, grazing significantly reduced soil OM (caged= 8.9 ± 0.4 %, uncaged= 7.5 ± 0.4 %; p =0.046; Table 1 and 2). This trend was similar in A3, though not significant (caged =14.1 ± 1.1, uncaged =13.4 ± 1.1 %; p =0.54; Table 1 and 2). Spring flush of TIN resulted in significantly higher concentrations (up to 3 times higher) in spring 2016 than all fall concentrations in A1 N (season x year p <0.0001; Table 1 and Table 2). This trend was similar in A3 in 2014 and 2015 only (season x year p =0.002; Table 1 and 2). There were no significant effects of grazing on TIN at either site, though in A1 N uncaged plots were slightly higher than caged plots (6.2 ± 1.8 versus 4.8 ± 0.6 mg/kg, respectively; p =0.09). In A1 N, there were no significant effects of either season or grazing on TP (Table 1 and 2); in A3, fall concentrations of TP were 1.2 times higher than spring (p <0.0001), and were negatively affected by grazing (p =0.02; Table 1 and 2).

**Table 1:** Avg ± SE soil characteristics measured in A1N and A3, throughout the study period. C= caged plots, U= uncaged control plots.

| Factor | Fall 2014 |  | Fall 2015 |  | Spring 2016 |  | Fall 2016 |  |
| --- | --- | --- | --- | --- | --- | --- | --- | --- |
|  | C | U | C | U | C | U | C | U |
| **A1N** |  |  |  |  |  |  |  |  |
| Organic matter (%) | 8.3 ± 0.9 | 6.9 ± 0.7 | 9.0 ± 1.0 | 7.2 ± 0.7 | 8.8 ± 0.5 | 7.2 ± 0.4 | 9.3 ± 1.0 | 8.6 ± 1.0 |
| Total inorganic nitrogen (mg/kg) | 4.0 ± 0.6 | 7.1 ± 3.5 | 1.5 ± 0.3 | 2.0 ± 0.5 | 11.3 ± 1.1 | 10.5 ± 1.1 | 2.4 ± 0.3 | 5.1 ± 1.9 |
| Total phosphorus (mg/kg) | --- | --- | --- | --- | 744.6 ± 23.5 | 683.7 ± 29.7 | 696.8 ± 22.3 | 724.9 ± 26.2 |
| **A3** |  |  |  |  |  |  |  |  |
| Organic matter (%) | 13.0 ± 0.9 | 12.5 ± 1.0 | 13.4 ± 1.1 | 12.5 ± 1.0 | 15.8 ± 1.2 | 14.6 ± 1.2 | 14.1 ± 1.0 | 13.8 ± 1.0 |
| Total inorganic nitrogen (mg/kg) | 15.4 ± 5.1 | 15.1 ± 3.3 | 2.6 ± 0.4 | 7.9 ± 3.7 | 25.6 ± 3.0 | 28.2 ± 5.4 | 12.9 ± 1.7 | 12.0 ± 1.4 |
| Total phosphorus (mg/kg) | --- | --- | --- | --- | 990.0 ± 53.2 | 903.0 ± 48.7 | 1176.2 ± 61.7 | 1102.3 ± 57.7 |

**Table 2:**
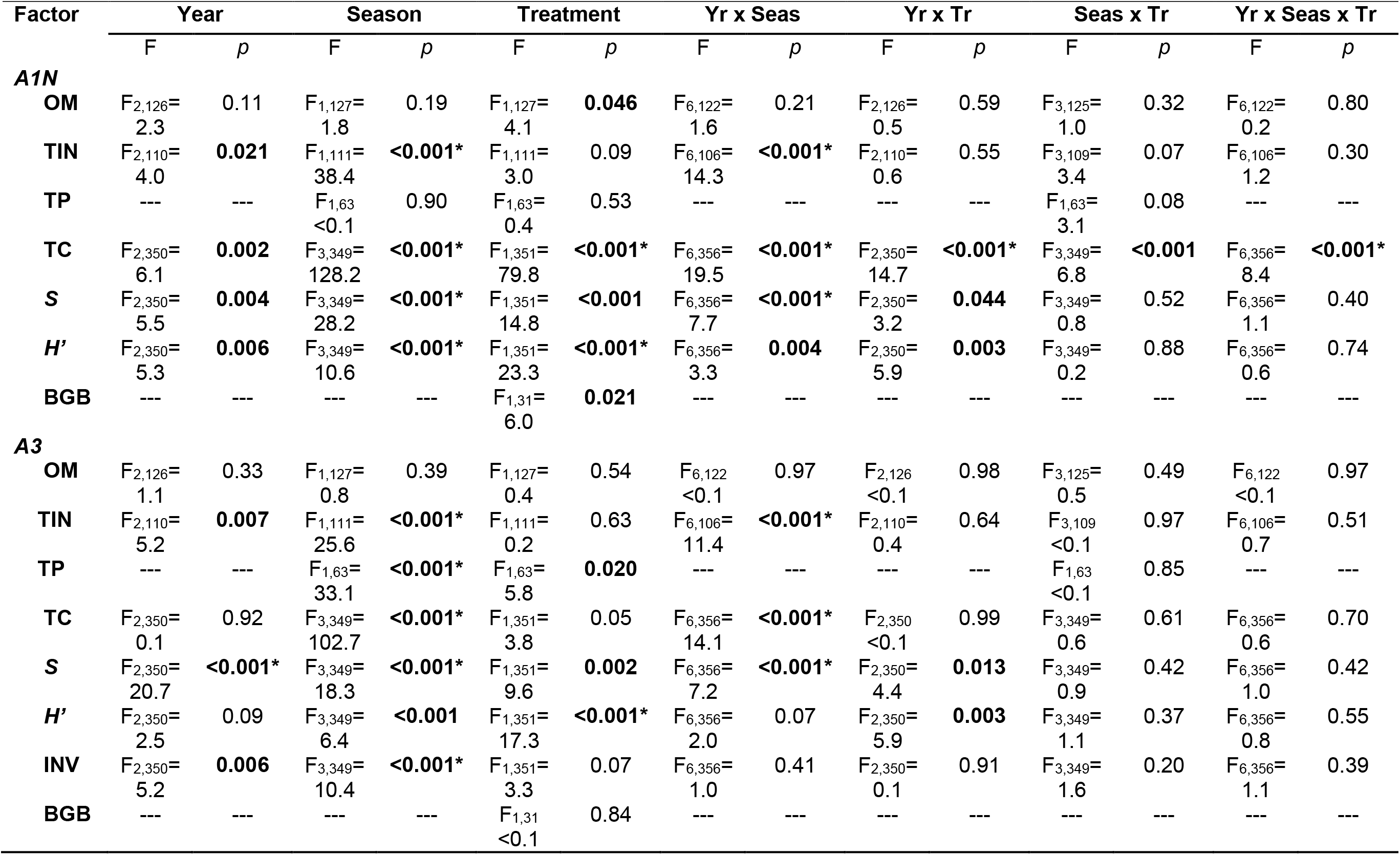
Results of analysis of variance examining the effect of year (2014-2016), season (spring, early summer, mid-summer, and fall), and grazing treatment (caged, uncaged) on environmental and plant characteristics at both wetland sites. One-way (belowground biomass [BGB]), two-way (soil total phosphorus [TP]), and three-way (% organic matter [OM], total extractable inorganic nitrogen [TIN], total plant cover [TC], plant species richness [S], and Shannon Wiener diversity index [*H’*] tests were used as appropriate. Significant p-values are bolded (*p <0.0001). Yr= year, Seas= season, Tr= treatment.

### Plant growth and diversity

Total plant cover was similar between A1 N and A3 in the control plots (p=0.11; Table S1). In A1N, grazing significantly reduced plant cover (Fig. 3), but a significant three-way interaction suggests that the impact of grazers varied by season and across years (p <0.0001; Table 2, Fig. 4a). The greatest grazing effect occurred in mid-summer (July), the height of the growing season (caged =112.7 ± 6.0, uncaged =81.8 ± 7.8 %). These effects increase over time, with the difference in cover between caged and uncaged plots (C-U) increasing from approximately 5% in 2014 to 55% in 2016. In A3, a significant two-way interaction also showed similar trends of total plant cover varying by season and across different years (p <0.0001; Table 2, Fig. 4b). Grazers slightly reduced plant cover in A3 (caged =61.4 ± 6.4, uncaged =55.1 ± 6.4 %; p =0.05). There were no differences between three-sided cage-control plots and uncaged control plots at either site (A1N: p= 0.56; A3: p =0.24). Belowground biomass in control plots was similar between A1N and A3 (260.2 ± 28.4 and 215.3 ± 39.1 g m^-2^, respectively; p =0.51; Table S1). In A1N only, grazing significantly reduced belowground biomass by 30% (A1N = 180.1 ± 17.7 g m^-2^, A3 = 204.7 ± 32.5 g m^-2^; p =0.021; Table 2).

**Fig. 3.**
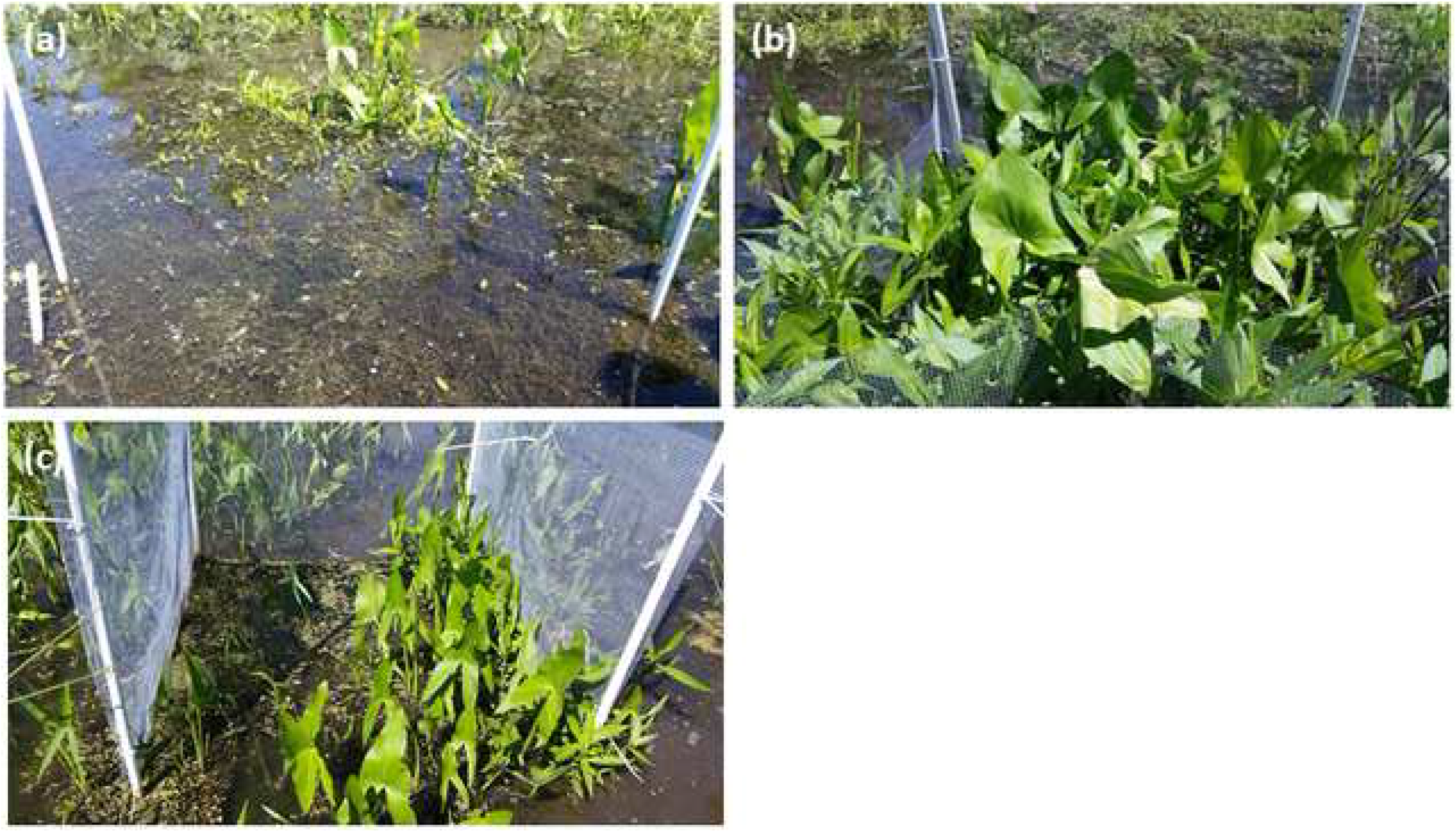
(a) uncaged control plot, (b) caged grazer exclusion plot and (c) three-sided cage-control plot during summer 2016.

Plant diversity was significantly lower in A1 N than A3 (*S* = 2.2 ± 0.3 and 3.9 ± 0.5, respectively, p <0.0001; *H’* = 0.4 ± 0.1 and 0.9 ± 0.1, respectively, p <0.0001; Table S1). In A1 N, the substantial reduction in diversity with grazing echoed total plant cover and increased over time for both *S* and *H’* such that caged plots had 1.3 and 1.7 times higher *S* and *H’*, respectively than uncaged plots in 2016 (p =0.044 and p =0.003, respectively; Table 2, Fig. 4c, 4e). Seasonal variation (peak in mid-summer) in *S* and *H’* also increased over time (*S*: p <0.0001; *H’*: p =0.004).

**Fig. 4.**
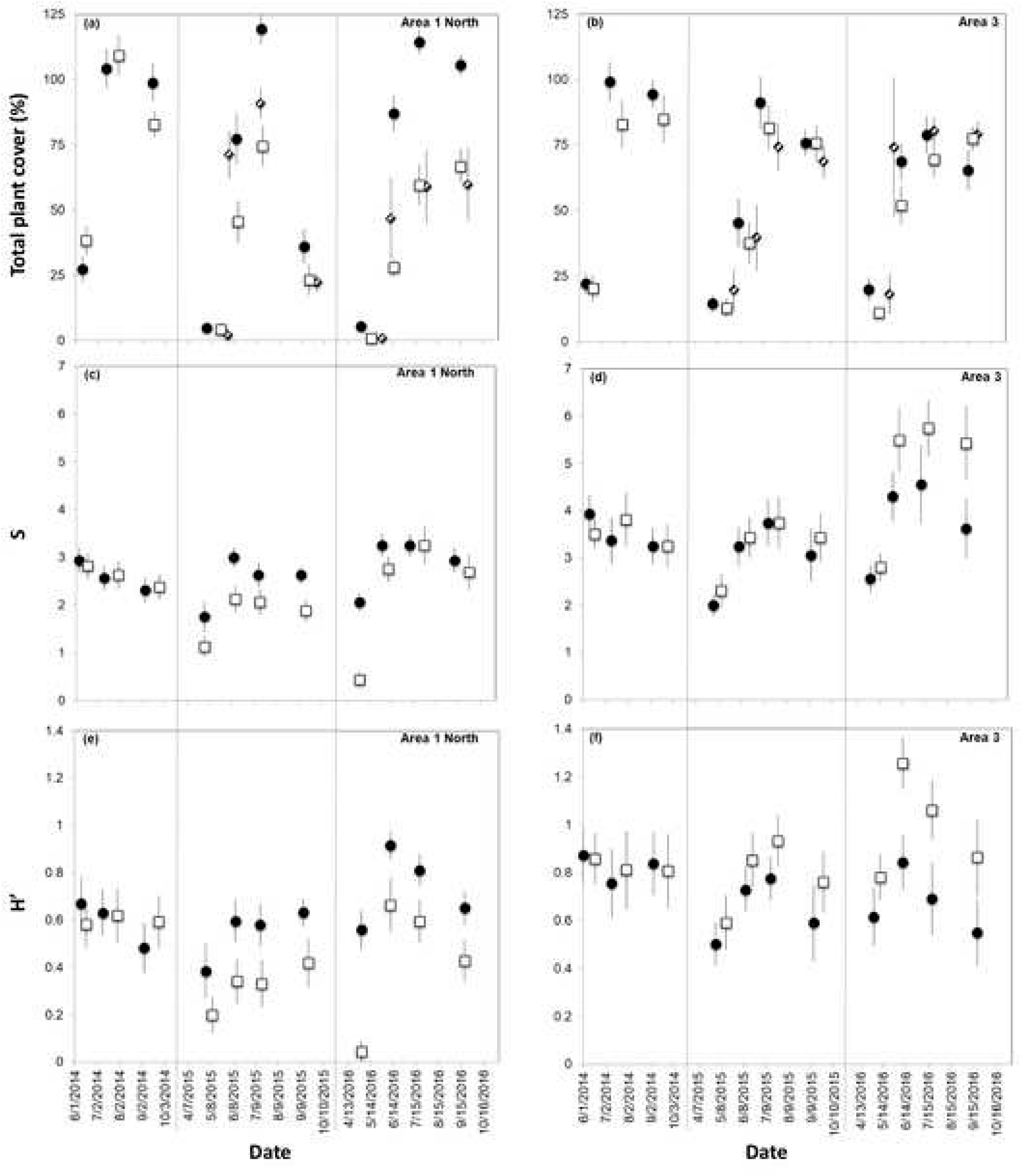
Mean ± SE plant characteristics measured in caged (black circle), uncaged (white square), and cage control (striped diamond) plots in study wetlands during the growing seasons of 2014-2016. Panels: (a) A1 N, total plant cover (b) A3, total plant cover; (c) A1N, species richness; (d) A3, species richness; (e) A1N, Shannon-Weiner diversity scores; (f) A3, Shannon Weiner diversity scores. Note that total cover may exceed 100% when plant canopies of individual species overlap.

In contrast to A1 N, grazing increased plant diversity in A3 and this effect, again, increased over time (*S*: p =0.013; *H’*: p =0.003; Table 2, Fig. 4d, 3f); in 2016, *S* and *H’* were 1.3 and 1.5 times higher in uncaged as opposed to caged plots (*S:* = 4.9 ± 0.6 and 3.8 ± 0.6, respectively; *H’*: 1.0 ± 0.1 and 0.7 ± 0.1, respectively). A3 also showed similar seasonal variations in diversity, which increased over time (*S*: p <0.0001; *H’*: p =0.07). Grazing reduced invasive cover in A3, but not significantly (caged =8.6 ± 3.3, uncaged =5.9 ± 2.3 %; p =0.07; Table 2, Fig. 5). Invasive cover was consistently highest in the fall (p <0.0001) and significantly decreased over the course of the study such that cover was 2.5 times higher in 2014 than in 2016 (10.3 ± 3.4 and 4.1 ± 1.9 %, respectively; p =0.006).

**Fig. 5.**
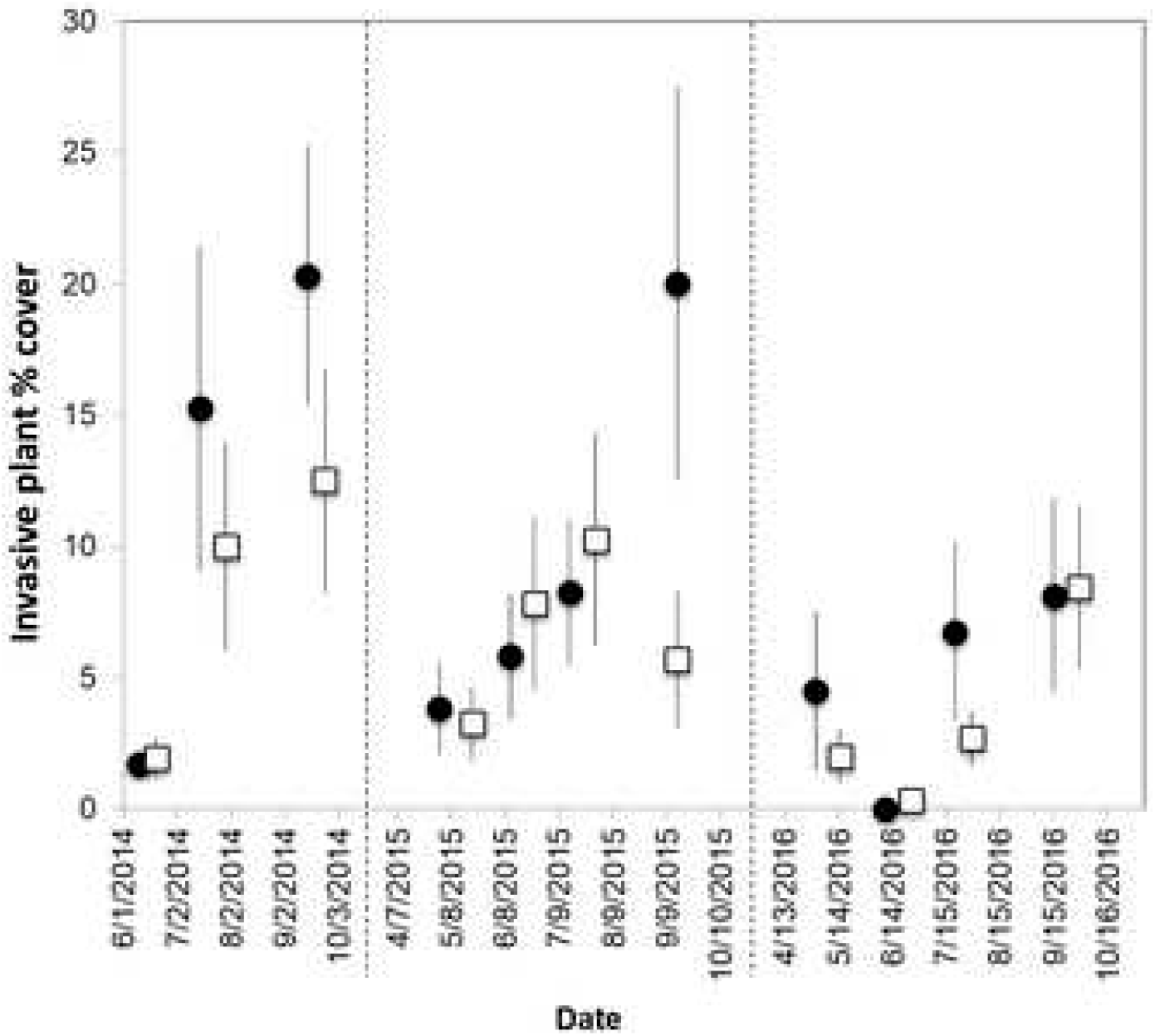
Mean ± SE invasive plant cover in caged (black circle) and uncaged (white square) plots in A3 wetland during the growing seasons of 2014-2016.

At both sites, stem height of the most common plant species was impacted more significantly by season and grazing treatment than stem density or individual species cover, but the trends were similar for all three variables (Fig. 6; Tables S2 and S3). At A1 N, where emergent wetland species dominated the community, the maximum effect of grazing coincided with the peak height and reduced stem height by 60-70% (*A. plantago-aquatica:* caged = 42.1 ± 6.8, uncaged = 16.6 ± 4.1 cm; *Polygonum* spp.: caged = 108.7 ± 6.7, uncaged = 29.6 ± 8.1 cm; treatment x season p <0.001 and p <0.0001, respectively; Fig. 6a, Table S2). Grazing also significantly reduced the height of *S. latifolia,* another dominant emergent species, by approximately 18%, though this was not seasonally dependent (caged = 73.1 ± 6.1, uncaged = 60.1 ± 5.1 cm; p = 0.041). In contrast, grazing significantly increased stem height for *Potamogeton amplifolius,* a submerged aquatic species, by approximately 35% (caged= 11.0 ± 0.3, uncaged= 17.1 ± 0.7 cm; p <0.0001). For other emergent species, *Leersia oryzoides* and *Schoenoplectus tabernaemontani,* the reduction in stem height was not significant (Fig. 6a, Table S2).

**Fig. 6.**
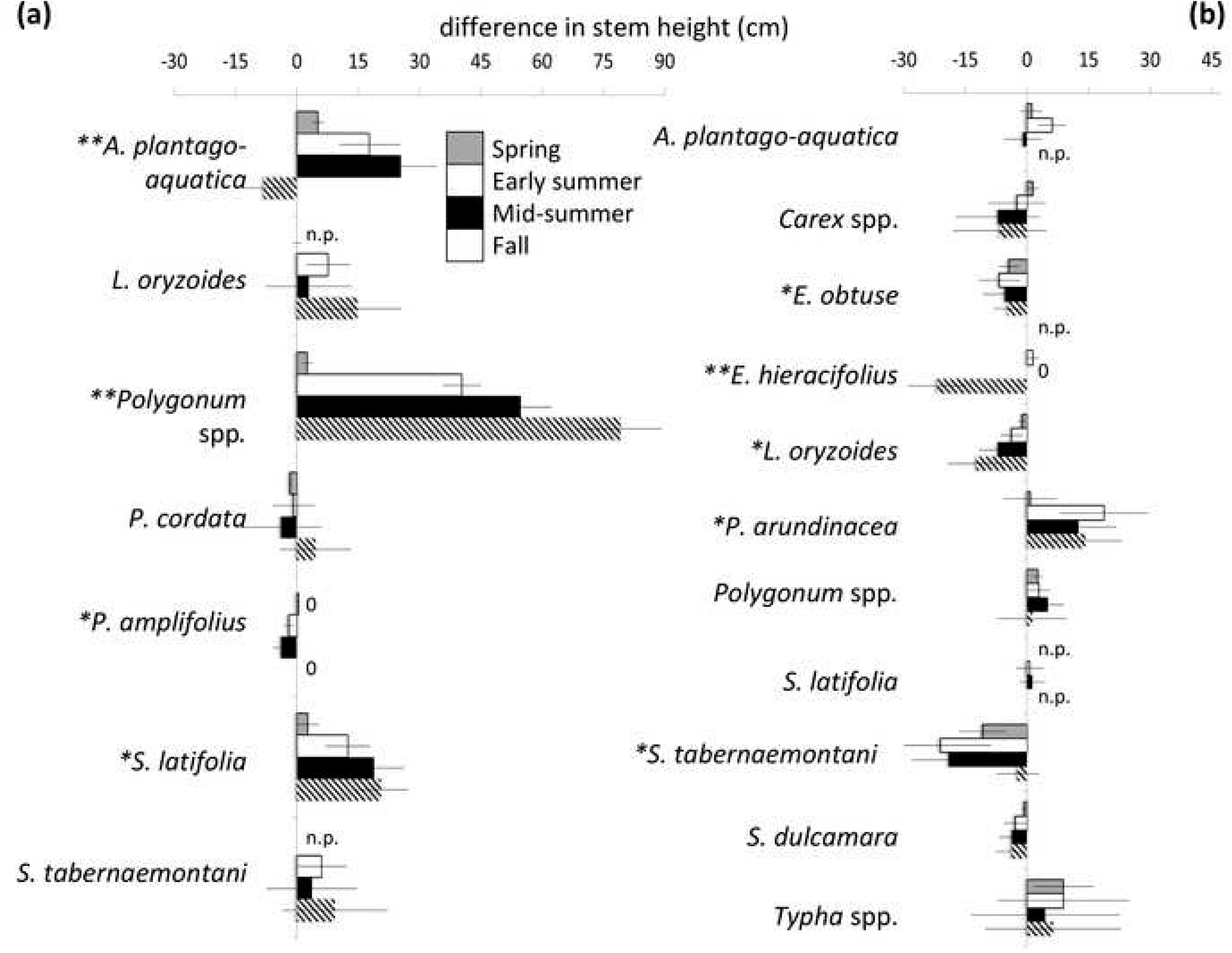
Mean ± SE difference in stem height (caged minus uncaged) of major species in A1N (a) and A3 (b) throughout the 2016 growing season. Positive values indicate caged > uncaged; negative values indicate uncaged > caged. For species where a value of 0 is indicated, caged = uncaged stem height and where n.p. is indicated, the species was not present in either plot. *grazing treatment p <0.05, **season x grazing p <0.05 based on two-way analysis of variance.

The plant community of A3 was characterized by a mixture of wet meadow, grasses, and emergent species. *Eleocharis obtusa, L. oryzoides,* and *S. tabernaemontani,* three native species, were significantly taller in ungrazed plots (p =0.048, p =0.02, p =0.004, respectively; Fig. 6b, Table S3). Stem height of *Erechtites hieracifolius,* another native species, was significantly greater in grazed plots, with a peak difference of 65% in fall (caged= 11.7 ± 5.8, uncaged= 33.5 ± 6.2 cm; treatment x season p <0.001). Conversely, grazing reduced stem height of *Phalaris arundinacea* and *Typha* spp., invasive species, though not necessarily to the same degree (F =5.6, p =0.02 and F =0.9, p =0.35 for height; Fig. 6b, Table S3).

## Discussion

Restoration to a stable community structure and function varies widely across ecosystems, depending strongly on antecedent conditions and management efforts. We found strong interactions between hydrology, nutrient availability, and herbivory that drove divergent and increasingly large responses to grazer exclusion in two created wetlands. In the permanently flooded wetland, the higher waterfowl grazing pressure coupled with lower nutrient availability led to substantial shifts in cover and diversity of emergent plants, promoting submerged species and a change in soil composition. With seasonal flooding and higher nutrient availability, herbivore access was lower and greater plant diversity and to some extent reduced invasive plant cover was observed in areas exposed to grazing. This suggests long-term consequences of intense grazing for habitat provision and delivery of other desirable ecosystem services.

Since hitting historic lows in the 1930s, waterfowl populations have increased due to extensive conservation efforts, widening of historic range limits with a changing climate, and increased agricultural land (Fox et al. 2005; Gauthier et al. 2005; Baldassarre et al. 2006). As numbers exceed limitations previously set by winter temperatures, vegetation losses up to 98% have been observed (Jefferies & Rockwell 2002; Gauthier et al. 2005). In this study, the ability of herbivores to influence wetland community dynamics was linked to temporal and spatial shifts in hydrology. Where higher water levels and open water permitted high densities of waterfowl (site A1 N), we found a significant reduction in both aboveground plant cover across the dominant species and an associated reduction in belowground biomass (Table 2, Fig. 3, 4a, 6). The greatest grazing impacts were observed in plots closest to a concealed goose thoroughfare between an adjacent pond and the created wetland. Spring flooding in A3 led to similar waterfowl abundance between sites, but with heterogeneous drying waterfowl abundance decreased. Our observations do not take into account grazing by larger insects, or nocturnal and crepuscular mammals, and while plant cover was impacted less in A3, the impact on plant diversity suggests a more complex influence of grazers on plant community structure.

Plant diversity at both sites (overall mean species richness in control plots, A1N = 2.2 ± 0.3 and A3 = 3.9 ± 0.5) was similar to other created and restored wetlands in the United States (2-6 species), but quite low when compared to natural reference wetlands in similar regions (10-12 species) (e.g., Brown and Bedford 1997; Campbell et al. 2002; Matthews et al. 2009). Grazer selectivity may lead to the removal of palatable species, lending a competitive advantage to plants with structural compounds or anti-herbivore phenolic compounds (Grosholz et al. 2009; Gutbrodt et al. 2012; Harrison et al. 2017; Levin et al. 2006; Morrison and Haye 2012). However, foraging effort is likely influenced by and varies depending on multiple factors including, but not limited to, predation risk, total food densities, and foraging location (Hagy and Kaminski 2015; Sherfy et al. 2011). In A1N, the substantial reduction of dominant, fast-growing species (*A. plantago-aquatica, Polygonum* spp., and *S. latifolia;* Fig. 6a, Table S2) suggests convenience rather than selectivity, and may explain the reduction in overall plant richness and diversity in response to grazing (Fig. 4c, 4e). Functional group shifts are subsequently reflected in the increased height, density, and cover of *P. amplifolius,* a submerged species, that perhaps flourished under the greater light availability in uncaged plots where large emergent plants were removed (Mitchell 1989; Shaffer et al. 2015).

Set in the different context of hydrology, nutrient availability, and grazing pressure, exclusion of grazers in A3 resulted in contrasting impacts on dominant species, suggesting dissimilar trajectories of wetland development. Grazer control resulted in neutral or positive trends in growth for many common native species, in contrast to the most common invasive species at the site *(P. arundinacea* and *Typha* spp.) that were negatively impacted. These species also exhibited significant leaf damage by vertebrates (deer clipping) and invertebrates (snail radulations) (Fig. 6b, Table S3). *P. arundinacea* and *Typha* spp., typically the tallest plants by early to midsummer, may have protected shorter native species from herbivore access (Bagousse-Pinguet et al. 2012; Barbosa et al. 2009). Low to moderate levels of grazing may lead to balanced competition among species, promoting greater overall survivorship and diversity (Hegland et al. 2013; Luo et al. 2012). These asymmetries between native and non-native plants in competitive ability and susceptibility to herbivores may explain their ability to coexist under moderate grazing pressure (Heard and Sax 2013). Without the mediating control of grazers, invasive species cover increased and diversity decreased as native species were out-competed (Fig. 4d, 4f, 5). This contrasts with the suggestion that invasive species have a competitive advantage because native grazers prefer native plants, or because invasive species contain novel chemical defenses intolerable for native grazers (Blossey & Notzold 1995; Callaway & Ridenour 2004). Invasive *P. arundinacea* was repeatedly introduced to North America as a forage grass for livestock, suggesting that it is both palatable and has some nutritional value and may be selectively grazed (Laverge and Molofsky 2004). The most common herbivorous waterfowl (*B. canadensis* and *Anas* spp.) observed at our two sites are generalist herbivores; their selection of plant species may be based on accessibility rather than palatability or nutritional quality. Such grazer selectivity (or lack thereof) is important to consider when seeding/planting created wetlands after construction in order to ensure persistence of desirable species.

The impact of grazer presence on overall height and dominance of plant species was greatest at the summer height of vegetation growth, though this did not correspond with the fall peak grazer abundance. Similar trends have been observed in natural aquatic systems, where high fall and overwintering waterfowl abundance has led to subsequent reductions in plant growth and distribution in the following growing season (Chaichana et al. 2011; Guillaume et al. 2014). The interaction between grazer intensity and timing, along with nutrient availability and species-specific physiologic response, determines whether specific plant species can compensate for the herbivory. In general, plants that are not limited by nutrients can compensate or respond positively to herbivory (e.g., Fornoni 2011; Proulx and Mazumder 1998). While it is challenging to directly link antecedent land use to present soil conditions and nutrient availability, gravel substrates, as found prior to restoration in A1 N, are often difficult to revegetate due to low nutrient-holding capacities (Hugron et al. 2011; Johnson 1987). The comparatively low soil nitrogen, phosphorus and organic matter, and low overall plant diversity at A1 N, a former gravel depository, even after several years, reflects these trends. In contrast, cattle deposit large quantities of nutrients into the soil through excretion of nutrient-rich feces (McGechan and Topp 2004; McGechan et al. 2008), as reflected in the substantially higher nutrients and organic matter found at A3. Grazing in the early stages of the growing season (April and May) may have less of an impact because of spring nutrient flushes, allowing for neutral growth compensation by the plants, when compared to periods of higher grazing intensity and lower apparent nutrient availability (June-September) (Fornoni 2011; Proulx and Mazumder 1998). While very high nutrient availability can lead to monocultures because some plants are released from limitation (Bobbink et al. 2010; Holdredge et al. 2010), under limiting conditions, herbivory may elicit different responses depending on timing of grazing pressure and nutrient availability.

In some systems, invasive plant removal alone can result in rapid recovery to desired structure (e.g., approximately 4 yr; Cuevos and Zalba 2010). Created wetlands that are seeded and/or planted with native species can demonstrate greater diversity over time, perhaps by reducing early colonization by invasive species (Balcombe et al. 2005; Collinge et al. 2011). However, wetland restoration projects frequently fail to produce successful results even after decades, with poor soil development as the most common shortcoming (Ballantine and Schneider 2009; Ballantine et al. 2014; Moreno-Mateos et al. 2012). Intense grazing in nutrient-poor emergent wetlands may be counter-productive to restoration efforts. Removal of aboveground material necessitates reallocation of stored below-ground resources towards recovery, substantially limiting – by 30% in A1N – the expansion of belowground root networks and rhizomes (Gao et al. 2008; Maron & Crone 2006; Piñeiro 2010). In the absence of grazing, higher plant biomass both above and belowground contributes to enhanced detritus and soil organic matter (Bai et al. 2012; Vaieretti et al. 2013). Failure to take into account the potentially transformational impacts of grazing on the trajectory of plant community development may contribute indirectly to poor soil development. Likewise, we suspect cascading impacts on biogeochemical cycling and delivery of services: as grazers continue to indirectly manipulate soil characteristics, wetlands become susceptible to reductions in carbon sequestration and increased emissions of greenhouse gases (Kayranli et al. 2010; Winton & Richardson 2016, Spangler 2019).

Created wetlands may, in part, be failing to meet functional and diversity expectations because young plant communities are vulnerable to shifts in composition initiated by herbivores (Moreno-Mateos et al. 2012; Seymour et al. 2010; Tanentzap et al. 2009), suggesting that herbivore control is necessary for plant community development (Sawtschuk et al. 2010). Increasing grazer impacts over time in this study suggest cumulative and interactive effects on plant communities with potential to shift the functional state from emergent to submerged vegetation where water is sufficiently deep, or from low to high diversity wet meadow species where it is not. As wetlands face more erratic rainfall patterns and summer drought conditions due to climate change, created wetlands that lack water storage capacity or control structures may be increasingly at risk. The creation of bathymetric heterogeneity within wetlands to include deep and shallow areas, stream rivulets, and microtopography by including distinct hydrological units and water control structures in wetland design will help to balance wetland use by grazer populations by providing separate areas for nesting and breeding, foraging, and open water that may be used seasonally (Ma et al. 2010; Yallop et al. 2004). This will also encourage the growth and development of multiple vegetation communities that may differ based on water depth, and provide different resources to herbivores based on varying diets, or habitat requirements and provide resilience in the face of extreme grazing or environmental fluctuations. Using small-scale protective enclosures to deter grazers, especially geese, initially after plantings will also help plant communities develop by protecting individuals during their most vulnerable growth period. Further study is required to understand these grazer-induced shifts in functional state and delivery of ecosystem services, such as carbon storage and nutrient removal.

## Acknowledgments

The authors gratefully acknowledge Waste Management of New York and the Thomas H. Gosnell School of Life Sciences and the College of Science at the Rochester Institute of Technology for supporting this work. We are grateful to Bruce Cady for participating in grazer monitoring, to Taylor Williams, Delanie Spangler, Kaitlyn Moranz, Elizabeth Bruen, Sonia Huang, Michael McGowan, Lisa Kratzer, Melissa Maurer Harrison, and Rachel Allen for assistance in the field, and to Nicole Fornof and Rebecca Zayatz for logistical support.

## Funding sources

The authors gratefully acknowledge Waste Management of New York and the Thomas H. Gosnell School of Life Sciences and the College of Science at the Rochester Institute of Technology for supporting this work

**Table S1:** Soil and plant characteristics in uncaged control plots (mean ± SE) and results of oneway analysis of variance comparing sites. Significant p-values are bolded and * indicates p <0.0001.

|  | A1N | A3 | F | p |
| --- | --- | --- | --- | --- |
| Organic matter (OM, %) | 7.5 ± 0.4 | 13.4 ± 0.5 | F<sub>1,111</sub>= 87.8 | **<0.001\*** |
| Total inorganic nitrogen (TIN, mg/kg) | 6.2 ± 1.8 | 17.1 ± 3.4 | F<sub>1,111</sub>= 17.1 | **<0.001\*** |
| Total phosphorus (TP, mg/kg) | 704.3 ± 28.0 | 1002.7 ± 53.2 | F<sub>1,63</sub>= 42.6 | **<0.001\*** |
| Total plant cover (TC, %) | 48.4 ± 5.5 | 55.1 ± 6.4 | F<sub>1,351</sub>= 2.5 | 0.11 |
| Species richness (S) | 2.2 ± 0.3 | 3.9 ± 0.5 | F<sub>1,351</sub>= 73.3 | **<0.001\*** |
| Shannon-Weiner (H') | 0.4 ± 0.1 | 0.9 ± 0.1 | F<sub>1,351</sub>= 74.4 | **<0.001\*** |
| Belowground biomass (BGB, g/m<sup>2</sup>) | 19.2 ± 1.9 | 21.8 ± 3.5 | F<sub>1,31</sub>= 0.4 | 0.51 |

**Table S2:** Results of two-way ANOVAs examining the effect of season (spring, early-summer, mid-summer, fall) and grazing treatment (caged/uncaged) on stem height (cm), stem density, and cover (%) for major plant species in Area 1 North. Minor species not analyzed: *Asclepias incarnate, Carex* spp., *Epilobium* spp., *Juncus effuses, Lythrum salicaria**, Nymphaea odorata, Sparganium americanum, Typha* spp.** Significant p-values are bolded (*p <0.0001; **invasive species). C= caged plots, U= uncaged control plots, Seas= season, Tr= treatment

| Factor | Avg ± SE |  | Season |  | Treatment |  | Seas x Tr |  |
| --- | --- | --- | --- | --- | --- | --- | --- | --- |
|  | C | U | F<sub>3,125</sub> | p | F<sub>1,127</sub> | p | F<sub>3,125</sub> | p |
| *A. plantago-aquatica* |  |  |  |  |  |  |  |  |
| Height | 19.6 ± 3.1 | 9.6 ± 1.7 | 15.98 | **<0.001\*** | 11.82 | **<0.001** | 6.32 | **<0.001** |
| Density | 5.0 ± 1.0 | 2.6 ± 0.6 | 3.30 | **0.023** | 4.58 | **0.034** | 1.04 | 0.376 |
| Cover | 4.6 ± 1.1 | 1.2 ± 0.2 | 4.75 | **0.004** | 10.22 | **0.002** | 2.57 | 0.058 |
| *L. oryzoides* |  |  |  |  |  |  |  |  |
| Height | 14.2 ± 4.1 | 7.8 ± 3.2 | 4.75 | **0.004** | 1.54 | 0.216 | 0.40 | 0.752 |
| Density | 4.4 ± 1.6 | 0.3 ± 0.1 | 1.15 | 0.332 | 6.58 | **0.012** | 0.87 | 0.459 |
| Cover | 3.5 ± 1.6 | 0.2 ± 0.1 | 1.18 | 0.322 | 4.53 | **0.036** | 0.82 | 0.485 |
| *Polygonum* spp. |  |  |  |  |  |  |  |  |
| Height | 62.3 ± 5.4 | 18.1 ± 3.0 | 84.25 | **<0.001\*** | 185.62 | **<0.001\*** | 24.39 | **<0.001** |
| Density | 21.0 ± 2.2 | 3.5 ± 0.8 | 6.44 | **<0.001** | 68.53 | **<0.001\*** | 2.10 | * |
| Cover | 21.9 ± 2.6 | 2.6 ± 0.6 | 18.84 | **<0.001\*** | 89.19 | **<0.001\*** | 13.60 | 0.104 |
|  |  |  |  |  |  |  |  | **<0.001** |
|  |  |  |  |  |  |  |  | * |
| *P. cordata* |  |  |  |  |  |  |  |  |
| Height | 7.8 ± 3.2 | 8.3 ± 2.8 | 1.85 | 0.142 | 0.01 | 0.915 | 0.18 | 0.908 |
| Density | 1.0 ± 0.4 | 1.7 ± 0.6 | 0.93 | 0.429 | 0.98 | 0.324 | 0.07 | 0.979 |
| Cover | 1.1 ± 0.5 | 2.9 ± 1.5 | 0.88 | 0.456 | 1.28 | 0.260 | 0.38 | 0.770 |
| *P. amplifolius* |  |  |  |  |  |  |  |  |
| Height | 0.5 ± 0.3 | 1.9 ± 0.7 | 4.37 | **0.006** | 4.28 | **0.041** | 2.22 | 0.089 |
| Density | 1.3 ± 0.8 | 2.0 ± 0.8 | 2.90 | **0.038** | 0.47 | 0.493 | 0.34 | 0.797 |
| Cover | 1.7 ± 1.0 | 1.4 ± 0.5 | 1.33 | 0.269 | 0.08 | 0.780 | 0.02 | 0.996 |
| *S. latifolia* |  |  |  |  |  |  |  |  |
| Height | 73.1 ± 6.1 | 60.1 ± 5.1 | 256.1 | **<0.001\*** | 18.84 | **<0.001\*** | 2.44 | 0.068 |
| Density | 17.0 ± 1.8 | 12.8 ± 1.2 | 3 | **<0.001\*** | 9.20 | **0.003** | 1.36 | 0.258 |
| Cover | 44.1 ± 4.6 | 29.9 ± 3.6 | 56.62 | **<0.001\*** | 16.27 | **<0.001\*** | 2.95 | **0.036** |
|  |  |  | 74.68 |  |  |  |  |  |
| *S. tabernaemontani* |  |  |  |  |  |  |  |  |
| Height | 8.4 ±3.8 | 2.9 ± 2.1 | 1.01 | 0.389 | 1.62 | 0.206 | 0.12 | 0.948 |
| Density | 1.1 ± 0.6 | 0.2 ± 0.1 | 0.27 | 0.847 | 2.15 | 0.145 | 0.29 | 0.835 |
| Cover | 0.8 ± 0.5 | 0.1 ± 0.0 | 0.81 | 0.809 | 2.72 | 0.102 | 0.45 | 0.720 |

**Table S3:** Results of two-way ANOVAs examining the effect of season (spring, early-summer, mid-summer, fall) and grazing treatment (caged/uncaged) on stem height (cm), stem density, and cover (%) for major plant species in Area 3. Minor species not analyzed: *Acer saccharum, Andropogon gerardii, Artemisia vulgaris**, Asclepias incarnate, Aster* spp., *Cornus sericea, Daucus carota, Echinochloa crus-galli, Epilobium* spp., *Juncus effuses, Juncus inflexus, Lactuca serriola, Lythrum salicaria**, Mimulus ringen, Ranunculus scelergtus, Rosa multiflora**, Rumex crispus*, *Solidago arguta*, *Sparganium americanum, Thinopyrum intermedium*, *Verbena hastate*. Significant p-values are bolded (*p <0.0001; **invasive species). C= caged plots, U= uncaged control plots, Seas= season, Tr= treatment.

| Factor | Avg ± SE |  | Season |  | Treatment |  | Seas x Tr |  |
| --- | --- | --- | --- | --- | --- | --- | --- | --- |
|  | C | U | F<sub>3,125</sub> | p | F<sub>1,127</sub> | p | F<sub>3,125</sub> | p |
| *A. plantago-aquatica* |  |  |  |  |  |  |  |  |
| Height | 14.0 ± 2.2 | 12.4 ± 1.8 | 23.21 | **<0.001\*** | 0.61 | 0.437 | 0.62 | 0.601 |
| Density | 4.7 ± 1.1 | 6.9 ± 1.5 | 13.88 | **<0.001\*** | 2.09 | 0.151 | 0.73 | 0.534 |
| Cover | 4.7 ± 1.2 | 3.7 ± 0.9 | 8.91 | **<0.001\*** | 0.58 | 0.447 | 1.04 | 0.378 |
| *Carex spp.* |  |  |  |  |  |  |  |  |
| Height | 7.8 ± 2.6 | 11.5 ± 3.7 | 3.28 | **0.023** | 0.74 | 0.391 | 0.22 | 0.886 |
| Density | 0.8 ± 0.3 | 1.7 ± 0.8 | 0.81 | 0.490 | 0.95 | 0.331 | 0.49 | 0.687 |
| Cover | 0.6 ± 0.2 | 0.6 ± 0.2 | 2.80 | **0.043** | 0.00 | 1.00 | 0.16 | 0.923 |
| *E. obtusa* |  |  |  |  |  |  |  |  |
| Height | 7.6 ± 2.1 | 12.9 ± 2.3 | 3.38 | **0.021** | 3.97 | **0.048** | 0.04 | 0.989 |
| Density | 2.5 ± 0.9 | 17.9 ± 5.3 | 3.00 | **0.033** | 9.78 | **0.002** | 1.92 | 0.130 |
| Cover | 0.4 ± 0.1 | 4.0 ± 1.4 | 2.07 | 0.109 | 7.52 | **0.007** | 1.27 | 0.288 |
| *E. hieracifolius* |  |  |  |  |  |  |  |  |
| Height | 3.3 ± 1.6 | 8.4 ± 2.4 | 28.97 | **<0.001\*** | 5.99 | **0.016** | 7.23 | **<0.001** |
| Density | 0.5 ± 0.4 | 1.6 ± 0.7 | 8.67 | **<0.001\*** | 2.28 | 0.134 | 2.37 | 0.074 |
| Cover | 0.5 ± 0.3 | 1.6 ± 0.6 | 13.86 | **<0.001\*** | 4.06 | 0.046 | 4.38 | **0.006** |
| *L. oryzoides* |  |  |  |  |  |  |  |  |
| Height | 3.0 ± 1.5 | 9.1 ± 2.4 | 4.25 | **0.007** | 5.00 | **0.027** | 0.78 | 0.507 |
| Density | 0.3 ± 0.2 | 4.1 ± 1.6 | 0.87 | 0.457 | 5.63 | **0.019** | 0.55 | 0.652 |
| Cover | 0.1 ± 0.0 | 3.3 ± 1.7 | 0.84 | 0.472 | 3.81 | 0.053 | 0.73 | 0.535 |
| *P. arundinacea*<sup>**</sup> |  |  |  |  |  |  |  |  |
| Height | 27.4 ± 6.1 | 15.8 ± 3.6 | 3.86 | **0.011** | 5.60 | **0.020** | 0.61 | 0.607 |
| Density | 8.5 ± 2.3 | 3.7 ± 0.9 | 0.79 | 0.504 | 6.22 | **0.014** | 0.38 | 0.765 |
| Cover | 10.6 ± 3.3 | 2.5 ± 0.7 | 0.18 | 0.908 | 7.65 | **0.007** | 0.09 | 0.964 |
| *Polygonum spp.* |  |  |  |  |  |  |  |  |
| Height | 27.0 ± 3.7 | 24.0 ± 3.3 | 57.09 | **<0.001\*** | 0.94 | 0.333 | 0.07 | 0.977 |
| Density | 27.4 ± 4.5 | 28.1 ± 4.7 | 15.95 | **<0.001\*** | 0.02 | 0.890 | 0.33 | 0.806 |
| Cover | 27.2 ± 4.3 | 25.2 ± 3.9 | 14.61 | **<0.001\*** | 0.24 | 0.626 | 2.58 | 0.057 |
| *S. latifolia* |  |  |  |  |  |  |  |  |
| Height | 3.5 ± 1.4 | 3.0 ± 1.1 | 5.90 | **<0.001** | 0.11 | 0.746 | 0.04 | 0.988 |
| Density | 2.4 ± 1.3 | 1.3 ± 0.6 | 2.86 | **0.040** | 0.75 | 0.387 | 0.26 | 0.858 |
| Cover | 2.1 ± 1.4 | 0.9 ± 0.4 | 1.85 | 0.142 | 0.77 | 0.383 | 0.29 | 0.834 |
| *S. tabernaemontani* |  |  |  |  |  |  |  |  |
| Height | 5.1 ± 2.6 | 18.4 ± 4.3 | 0.45 | 0.718 | 8.43 | **0.004** | 0.88 | 0.455 |
| Density | 0.4 ± 0.2 | 3.9 ± 1.1 | 0.79 | 0.501 | 12.51 | **<0.001** | 1.05 | 0.373 |
| Cover | 0.2 ± 0.1 | 1.9 ± 0.7 | 0.86 | 0.465 | 7.54 | **0.007** | 0.74 | 0.528 |
| *S. dulcamara*<sup>**</sup> |  |  |  |  |  |  |  |  |
| Height | 0.8 ± 0.7 | 3.6 ± 1.6 | 0.94 | 0.424 | 2.61 | 0.109 | 0.19 | 0.903 |
| Density | 0.1 ± 0.1 | 0.3 ± 0.2 | 9.43 | **<0.001\*** | 2.48 | 0.118 | 2.57 | 0.057 |
| Cover | 0.1 ± 0.1 | 0.5 ± 0.2 | 0.43 | 0.734 | 3.19 | 0.077 | 0.05 | 0.987 |
| *Typha spp.*<sup>**</sup> |  |  |  |  |  |  |  |  |
| Height | 39.6 ± 7.6 | 32.5 ± 6.4 | 5.98 | **<0.001** | 0.89 | 0.348 | 0.02 | 0.996 |
| Density | 1.3 ± 0.3 | 1.2 ± 0.3 | 3.27 | **0.024** | 0.13 | 0.717 | 0.01 | 0.999 |
| Cover | 2.0 ± 0.4 | 2.6 ± 0.7 | 2.47 | 0.065 | 0.85 | 0.357 | 0.39 | 0.757 |

